# Measurement of junctional tension in epithelial cells at the onset of primitive streak formation in the chick embryo via non-destructive optical manipulation

**DOI:** 10.1101/501775

**Authors:** Valentina Ferro, Manli Chuai, David McGloin, Cornelis Weijer

**Affiliations:** Dept of Physics, School of Science and Engineering, University of Dundee, Dundee DD1 4HN, UK.; School of Life Sciences, University of Dundee, Dundee, DD1 5EH, UK.; School of Electrical and Data Engineering, University of Technology Sydney, Ultimo, NSW 2007 Australia

## Abstract

Oriented cell intercalations and cell shape changes are key determinants of large-scale epithelial cell sheet deformations occurring during gastrulation in many organisms. In several cases directional intercalation and cell shape changes have been shown to be associated with a planar cell polarity in the organisation of the actin-myosin cytoskeleton of epithelial cells. This polarised cytoskeletal organisation has been postulated to reflect the directional tension necessary to drive and orient directional cell intercalations. We have now further characterised and applied a recently introduced non-destructive optical manipulation technique to measure the tension in individual cell junctions in the epiblast of chick embryos in the early stages of primitive streak formation. We have measured junctional tension as a function of position and orientation. Junctional tension of mesendoderm cells, the tissue that drives the formation of the streak, is higher than tension of junctions of cells in other parts of the epiblast. Furthermore, in the mesendoderm junctional tension is higher in the direction of intercalation. The data are fitted best with a Maxwell model and we find that both junctional tension and relaxation time are dependent on myosin activity.

## Introduction

Development is characterised by the formation and shaping of new tissues of increasing complexity. An early critical phase of tissue formation is gastrulation where the main three germlayers, the ectoderm, mesoderm and endoderm, are formed and take up their correct topological positions in the embryo ^1^. In amniote embryos gastrulation starts with the formation of the primitive streak, the structure through which the mesoderm and endoderm precursors move into the embryo ^2, 3^ The process of streak formation is widely studied in chick embryos, since these can be cultivated in vitro and are accessible to manipulation ^4^. The embryo derives from a single layer of epithelial cells, the epiblast; the streak starts to form in the posterior part of the epiblast and extends in the anterior direction during its formation. Streak formation has been shown to involve large scale vortex like tissue flows in the epiblast ^5, 6, 7, 8, 9^. The vortex flows initiate in the sickle shaped area of the posterior epiblast gives rise to the endoderm and mesoderm (Fig 1A). Using a recently developed transgenic chick strain in which the cell membranes are labelled with GFP and the development of a dedicated lightsheet microscope we have previously been able to observe the process of streak formation at both the tissue and cellular level in considerable detail ^10^.The cellular mechanisms that have been proposed to drive these flows involve directed cell shape changes and cell intercalations and are supported by cell divisions and ingression of individual cells in the epiblast ^10, 11^. Before the onset of the tissue flows, the mesendoderm precursor cells are elongated and aligned in the direction of the forming streak. The onset of motion is correlated with cell shape changes and cell intercalations perpendicular to the A-P axis in the mesendoderm. Aligned cells form transient chains of junctions and these junctions are enriched in active myosin as detected by phosphorylation of the myosin light chain ^10^ (Fig. 3A). Blocking of myosin II activity relaxes cell shapes and inhibits directional cell intercalations and streak formation. Further experiments showed that blocking of myosin I resulted in a relaxation of the cells and absence of the formation of myosin II cables in aligned cell junctions ^10^.

**Figure 1:**
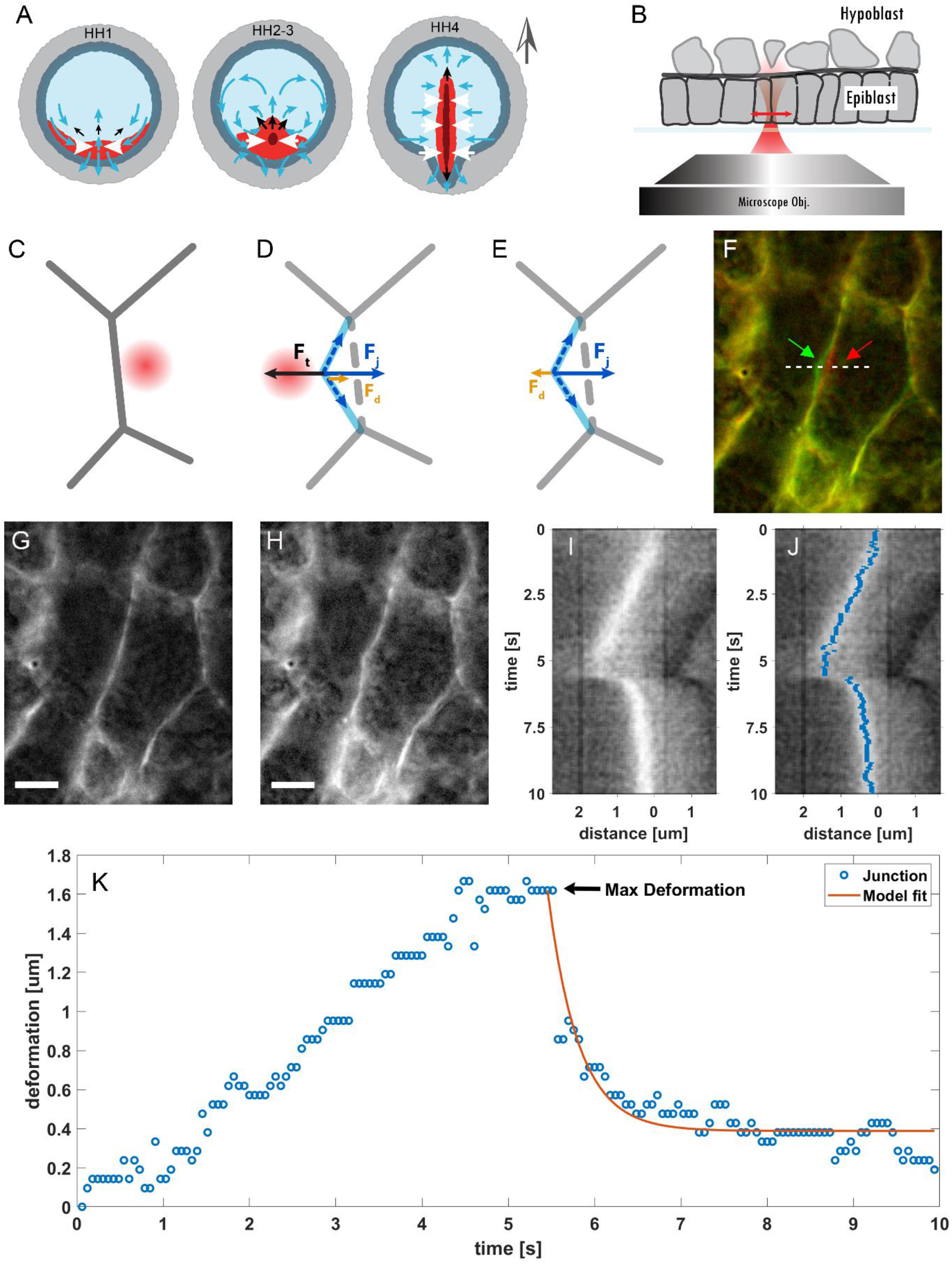
Optical manipulation of cell-cell junctions –. A) Stages 1-4 of the chick embryo development according to Hamburger-Hamilton ^35^. The different regions in the embryo are shown with different colours. The central part of the embryo, known as the *area pellucida* (light blue region) will form the embryo proper and is separated from the extra embryonic region the *area opaca* (grey region) by the marginal zone (dark green region). The presumptive mesendoderm (red region) is located in the posterior area of the embryo next to the marginal zone and will form the streak. At stage HH1 the mesendoderm cells start to move (blue arrows) due to active pulling forces (white arrows) generated in this tissue. The contraction of this tissue generates pushing forces (black arrows) that result in elongation of the streak at stages HH2-3. From stage HH3 onwards the mesendoderm cells start to ingress into the embryo through the streak. The grey arrow outside the embryo indicates the A-P axis. B) Schematic of the experiment in a cross sectional view: the chick embryo is situated on a glass-bottom plate with the epiblast facing the microscope objective. The optical trap is moved perpendicular to the cell-cell junctions (double red arrow). C-E) Bottom view of the experiment: the trap is turned on while on the right side of a selected junction (C) and then moved across the junction, once the trap crosses the junction, it deflects it (D). The maximum deformation is obtained when the optical force Ft is balanced by the tension of the junction Fj and the drag in the cytosol Fd. When the trap is turned off (E), Fj restores the junction to its rest position. F-H) False colour image corresponding to two time frames (F): red channel is the junction at rest position at t=0 (G), green channel is the junction at its maximum deformation (H). The images are extracted from Movie 1. Scale bar indicates 5μm. I) Kymograph of the junction deformation collected at the row indicated by the white dashed line in (G). J) Superposition of kymograph in (G) and the junction position extracted by the seam carving algorithm (blue pixels). K) Junction position as extracted from the kymograph and corresponding fit using the viscoelastic model.

**Figure 3:**
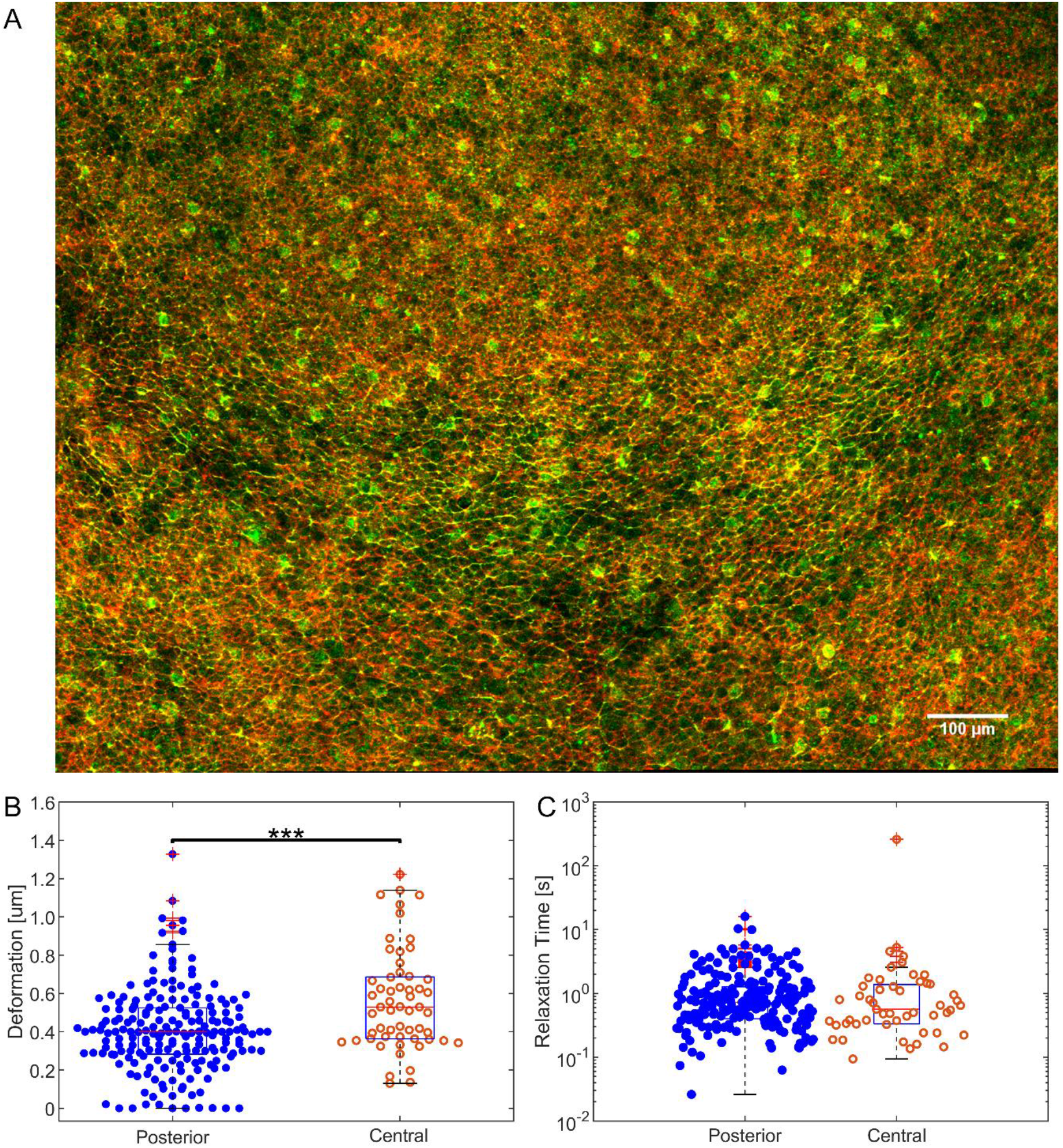
Junction deformation and relaxation time for the posterior and the central area of the embryo. (A) Image of posterior part of embryonic area of a chick embryo, posterior is to the bottom, anterior is to the top. The image shows phosphor specific myosin light chain antibody staining in green and actin detected by phalloidin staining in red. Note the organisation of supercellular phospho-myosin cables arranged in a horizontal arc in the mesendoderm domain located in the middle slice of the embryo. White scale bar, 100μm. (B) Boxplot and distribution of the maximum deformation of junctions measured in the posterior (blue dots) and in the central areaof the embryo (orange circles). (C) Boxplot and distribution of the relaxation times. In the posterior area, a median deformation of 0.39μm was measured, while in the central area the median deformation was 0.58μm. The measured relaxation times for this data set are 0.7 s and 0.6 s for the posterior and the central area respectively. The data for the posterior area were aggregated from ten different embryos, the same referred to as “Control” in Fig. 4 (n=203). The data for the central area were aggregated from three different embryos (n=57). *** indicates a p-value <0.001.

**Figure 4:**
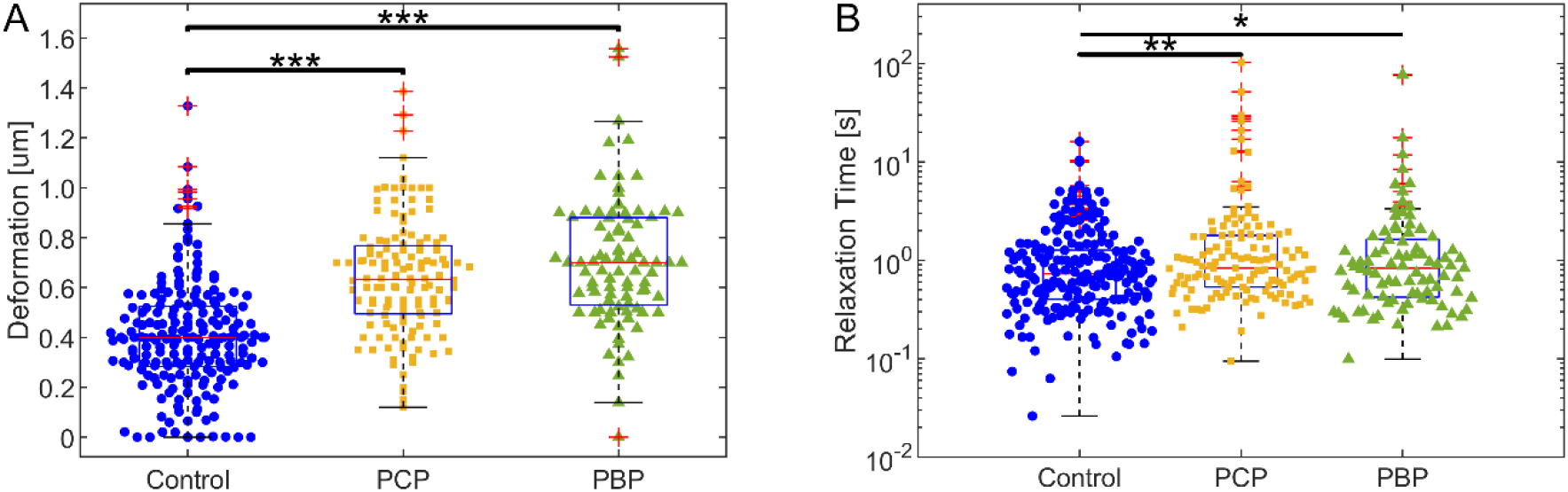
Junction deformation and relaxation time for embryos treated with myosin inhibitors. Boxplot and distribution of the maximum deformation (A) and of the relaxation time (B) of junctions measured in the posterior embryos before (blue dots, n=203) and after being treated with PCP (yellow squares, n=132) or PBP (green triangles, n=88). Median values for the deformations are 0.39μm, 0.62μm and 0.71 μm for the control, the PCP and the PBP data sets respectively. Median values for the relaxation times are 0.7s for the control sample and 1s for both the PCP and PBP treated samples. The control data were aggregated from ten different embryos, while the data for PBP and PCP are aggregated from three different embryos each. * indicates p-values <0.05; ** indicates p-values <0.01; *** indicates p-values < 0.001.

The cellular behaviours observed during gastrulation in the chick embryo thus appear to show very strong similarities to those that drive gastrulation in Drosophila, a process that involves extension of the body axis and is known as germband extension. Germband extension has been shown to depend on directional cell intercalations, driven by myosin II mediated directional shorting of junctions ^12, 13, 14^. It has furthermore been shown by localised laser ablation experiments that the contracting junctions are under greater tension than non-contracting junctions ^15^. Recently a more quantitative method has been developed to measure the tension in individual junctions directly by pulling on them, using a laser tweezers approach^16^. From the extension of the junction and the kinetics of a junction as it returns to the resting state after release from the trap, it was possible to extract parameters such as stiffness and viscosity of the junction. It was shown that the stiffness is around 2.5 fold higher in contracting junctions than in parallel non-contracting junctions during germband extension, while the measured viscosity can account for long term dissipation of tension ^17^. The latter was shown to be dependent on remodelling of the actin cytoskeleton.

Our working hypothesis to explain gastrulation in the chick embryos is that myosin II activity drives cell shape changes and importantly directional intercalation through the generation of directional tension. Testing this hypothesis requires measurement of junctional tension under a variety of experimental conditions. Here we demonstrate a modified version of the optical trapping technique used to measure tension in Drosophila junctions in [16] and find that we can optically manipulate junctions in cells of the epiblast of early gastrulation stage chick embryos. We observed, however, that in most cases we did not trap the junctions, but small vesicles in epiblast cells. These vesicles are then used as probes to deform the membrane. We performed relative tension measurements in early streak stage embryos and found differences in junctional tension in different areas of the embryo and, through the use of specific inhibitors, have defined a contribution of Myosin I and Myosin II in generating tension in these junctions.

## Results

Mechanics, especially the tension of cell junctions, are a major determinant of cell shape which in turn underlies tissue shape^18^. Changes in junctional tension drive a diverse array of cell behaviours. The mechanical properties of cell-cell junctions can be determined by applying external forces to the junctions while measuring their responses to these perturbations. We opted for optical tweezers to perform tension measurement in chick embryos, as this allowed us to apply optical forces to probe the system in a non-destructive manner, contrary to commonly used methods such as laser ablation, and without the need to introduce external probes, such as oil droplets in the embryos^19, 20^. We can measure the dynamics of a junction after it is moved a given distance from its equilibrium position. The deformation, which results in increased tension in the junction, generates a restoring force. When the junction is released, the tension enables the junction to contract back to its rest position^16^. To monitor the deflection of the junction, we used a combination of transgenic embryos expressing a membrane-localized GFP and high resolution fluorescence microscopy.

We designed and built an instrument integrating optical tweezers in an inverted fluorescence microscope ^21^. We used a single microscope objective to both focus the light from a high intensity infrared laser to produce the tweezers and to illuminate the sample with excitation light suitable to excite the GFP in the cell membranes (Fig. S1).

In our experiments, the trapping laser sweeps at right angles over the junction to be trapped in a single pass, starting in a cell on one side of the junction and ending up across the junction in a neighbouring cell, after which the trap is turned off (Fig. 1B-E). We used 750mW of laser power (measured in the image plane) and a measurement cycle of 10 seconds. This gave a good compromise between a small but measurable deflection of the majority of the tested junctions and absence of any obvious visible signs of damage (Fig. S2). We observed that when the trap is activated, the optical force overcomes the other forces acting on the junction (i.e. the tension and the drag from the cytosol) and it starts to deform (Fig. 1D and Fig. 1F-H). The junction reaches a maximum deformation when all the forces acting on it are in equilibrium (Fig. 1C-E and Fig. 1F-H). When we turn the trapping laser off, the junctional tension restores the junction to its rest state, opposed only by the drag in the cytosol (Fig. 1E). Stiffer junctions are lost from the moving trap before it reaches its final position.

To study the deformation over time, we generated kymographs of the junctions (Fig. 1I). The kymograph shows that the junction follows the tweezers until it reaches its maximum deformation. When the trap is tuned off, the junction then comes back to, or close to, its original position. This return phase follows an exponential decay determined by a time constant characteristic of the mechanical properties of the junction and its surroundings.

To perform the analysis, we extracted the position of the junction over time by applying a seam carving algorithm to the kymographs (Fig. 1J-K). This method is limited to pixel resolution, but we observed that in our experiments it outperformed other more typical approaches, such as a Gaussian fit of the image to localise the junction (Fig. S3). We then fitted the junction deformation over time and extracted the maximum deformation and the relaxation time.

### Optical trapping of cell-cell junctions

The optical tweezers used to optically manipulate of cell-cell membranes in living embryos (Drosophila) were calibrated by assuming that they would directly interact with and trap the cell-cell junction ^16, 17^. However, during our experiments in the chick embryo, we noticed that in most cases small vesicular organelles in the proximity of junctions were closely following the movement of trap (Fig. 2). By using a Fourier bandpass filter, we enhanced the contrast of these objects and observed how the organelles trapped by the tweezers ultimately are responsible for pushing and deforming the junctions (Fig. 2).

**Figure 2:**
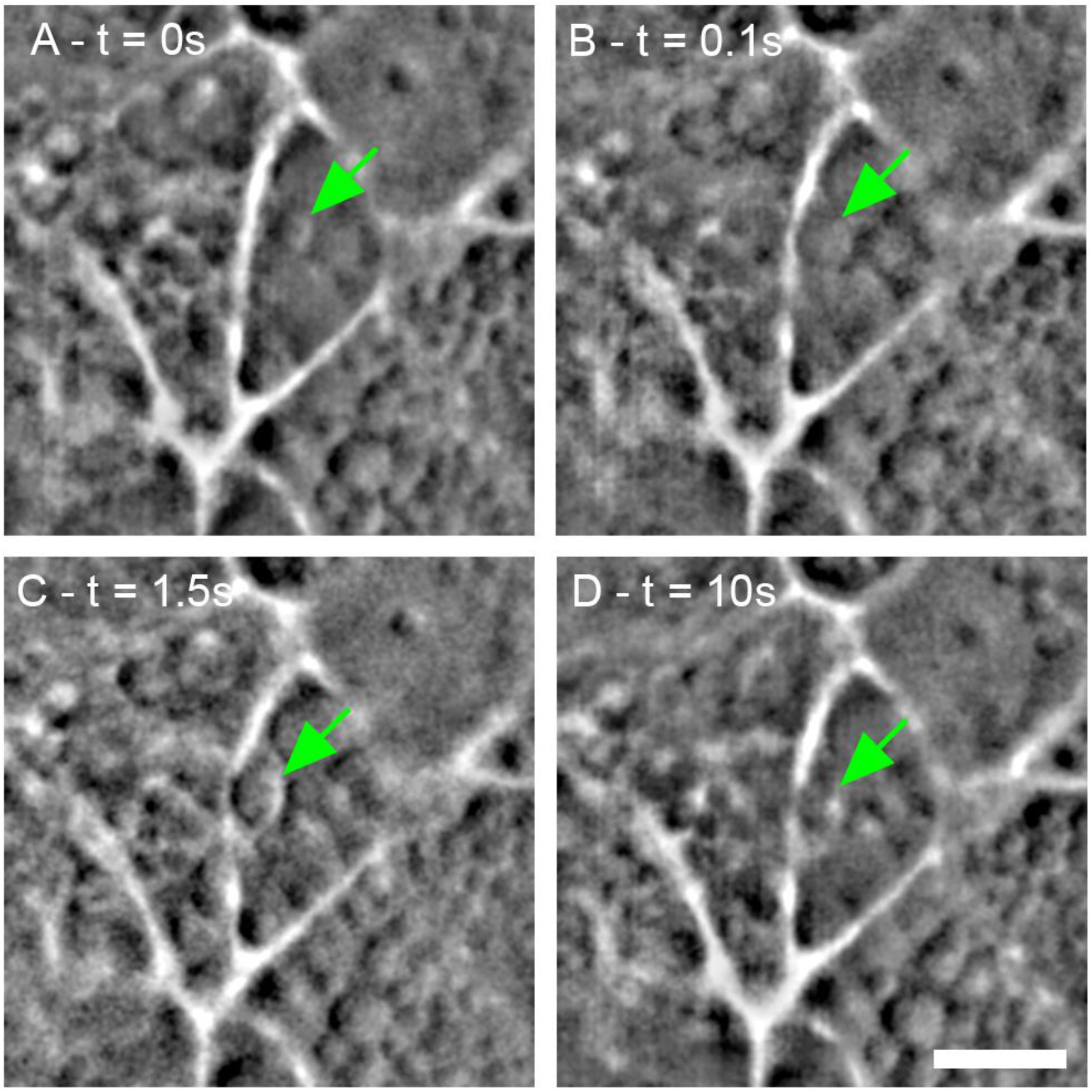
Contribution of organelles in the deformation of the junctions –. Sequence of pull & release experiment figures where an organelle (green arrow) is trapped and drives the deformation of the junction. The sequence is extracted from Movie 2. (A) At time t=0, the trapping laser is off, and an organelle is visible in proximity of the junction. (B) Once the trap is turned on, t=0.1s, the organelle is attracted into the trap. (C) With the trap moving, the organelle pushes against the junction until it reaches the maximum deformation at t=1.5s. (D) After switching the trap off, the junction and organelle return to their rest positions, with the organelle adjacent on the junction’s right side (t=10s). Scale bar indicates 5μm.

The observation that in most cases we likely trap organelles that then push the junction plays an important role in the interpretation of the results. The stiffness of optical tweezers is depended on the refractive index, the shape and the size of the trapped object ^22 23^. Since the trapped vesicles vary in size, the stiffness of the tweezers will vary also between measurements making generic calibration much more difficult.

Even though the trapping of vesicles resulting in varying stiffness of the trap introduces scatter in the data, we could still compare the response of junctions under different experimental perturbations and in different places in the embryo. Maintaining the properties of the laser trap such as laser power, distance travelled by the tweezers and duration of the experiment constant, we expect that higher tensions will result in smaller maximum deformation of the junctions. We implemented a constant perturbation protocol and measured the responses of the junctions in different parts of the embryo and under different experimental perturbations. We measured 40-100 junctions per condition and measurements were derived from 3-5 embryos of similar stages of development.

### High junctional tension in mesendoderm cells

The accumulation of myosin in junctions of mesendoderm cells is assumed to generate a significant junctional tension that will drive the directed cell-cell intercalations that mediate the elongation of the streak (Fig 3A). We wanted to verify the hypothesis that junctions oriented perpendicular to Anterior-Posterior (A-P) axis in the posterior mesendoderm precursor cells in the epiblast of embryos about to form a streak are under higher tension compared to junctions elsewhere in the embryo. If this were the case, we would expect that mechanical perturbation of these junctions would result in a smaller deflection of these junctions compared to junctions elsewhere in the embryo.

We therefore measured deformation in junctions aligned perpendicular to the A/P axis in the posterior mesendoderm part of the embryo and compared this with the deformation of junctions in the central part of the embryo before the formation of the streak. We observed a significant difference in the maximum deformation measured (Fig. 3B). The maximum deformation measured in the posterior area was 0.39 μm, (median, n=203,90% Confidence Interval CI [0.13 μm, 0.65 μm]), while in junctions located outside the mesendoderm in the central area of the embryo we measured a value of 0.58μm (median, n=57, 90% CI [0.32 μm, 0.99 μm]). Fitting the junctions with a Maxwell viscoelastic model, we extracted the relaxation time for each junction. The difference between the two distributions was not significant, with median values of 0.7s (n=203, 90% CI [0.3s, 3.1s]) and 0.6s (n=57, 90% CI [0.2s, 2.8s]) for junctions in the posterior and the central area respectively (Fig. 3C). Thus, the observed differences in junction deflection amplitude of cells located in the central and the posterior areas of the embryos suggest that the junctional tensions in these areas are different. In a different experiment where we used a sinusoidal movement of the optical trap across the junction, we found that junctions in Koller’s Sickle oriented along the A-P axis, were deformed more easily than junctions aligned perpendicular to the A-P axis, i.e. junctions aligned along the direction of the myosin cables (Fig. S4). Additional results show that junctions of cells in very young embryos, incubated for only 3 hours and optically manipulated before the formation of Koller’s Sickle, were more easily deformed than the junctions of embryos manifesting a clear Koller’s Sickle (Fig.S5). These measurements provide further support to the hypothesis that differences in tension drive differences in cell behaviours such as directional cell intercalation.

### Effect of Myosin inhibitors

Our previous work has shown that myosin II plays a key role in the execution of direction cell intercalation and streak formation ^10^. Both inhibitors of myosin II and siRNA mediated down regulation of myosin IIa and myosin IIb resulted in inhibition of directional cell intercalation and streak formation. We also found that inhibition of myosin I family members through inhibitors or specific siRNAs also resulted in a strong reduction of myosin II activity, as measured by inhibition of the formation of cables of myosin light chain phosphorylation, which resulted in a complete inhibition of directional cell intercalation and a block of streak formation. To investigate the roles of myosin I and myosin II in the generation of junctional tension in the mesendoderm cells we inhibited the activity of these myosins using specific inhibitors. We investigated the effects of two Myosin inhibitors: pentachloropseudilin (PCP) which specifically inhibits members of the myosin I family, and pentabromopseudilin (PBP), that is known to inhibit myosin II ^10, 24, 25^. We used a concentration of 10 μmolar for both inhibitors.

We measured the response of cell-cell junctions perpendicular to the A-P axis in the posterior area of embryos before and after the embryos were treated with these inhibitors. We measured a significantly larger junctional deformation after the embryos were treated with both myosin I and myosin II inhibitors. After the treatment with PBP, the maximum deformation almost doubled from 0.39μm (median, n=203, 90% CI [0.13 μm, 0.65 μm]) in the control data to 0.71μm (median, n=88, 90% CI [0.44 μm, 0.99 μm]) after inhibitor treatment. The treatment with PCP also caused the maximum deformation to increase to reach 0.62μm (median, n=132, 90% CI [0.35 μm, 0.95 μm]), results that were statistically highly significant, as per the results of t-tests performed. These experiments show that myosin II activity is a major determinant of junctional tension. They furthermore show that inhibition of myosin I activity results in a large decrease in junctional tension, consistent with its reported effect on myosin II cable formation. Finally, when we extracted the relaxation time from the fitting of the junctions, we observed a significant difference between the distributions when performing t-tests on the distributions, with a median of 1s for the PCP (median, n=132, 90% CI [0.4s, 3.1s]) and PBP (median, n=88, 90% CI [0.3s, 2.3s]) case compared to the 0.7s measured in the case of control junctions. The fact that the data in all experiments were on aggregate best fitted with a Maxwell viscoelastic model suggested that the duration of the perturbation might affect the irreversibility of the junction deformation (Fig. S7). This was clearly demonstrated in recently reported measurements of the effect of perturbation time on the irreversibility of the junctional deformation in Drosophila ^17^. This prompted us to see whether this effect was noticeable in our measurements. As explained before, under our conditions junctions are lost from the trap at various times depending on trap stiffness and junction tension. We therefore analysed the irreversibility obtained from the data fit as a function of perturbation time. The results show that up to 5 seconds the duration of the perturbation the effect was very small. However, after application of the PCP and PCB inhibitors, the irreversible viscous behaviour became more noticeable after shorter perturbation times (Fig S6B), suggesting that myosin activity may lower the viscous time constant of the junctions.

In conclusion these measurements show that myosin I and myosin II contribute significantly to junctional tension in mesendoderm cells and that tension is higher in mesendoderm cells junctions that are aligned perpendicular to the A/P axis along the direction of contraction.

## Discussion

While optical tweezers are an established tool for molecular biology and cellular biology, only recently they have been used in living organism, such as Drosophila embryos ^16^ and zebrafish ^26 27^. The application of optical tweezers for the manipulation of cell junctions in living organism was first reported by Lenne and co-workers where it was applied to measure the tension in junctions in the Drosophila embryo ^16^. Our experiments in the chick embryo have shown that these optical tension measurements are possible in more complex systems. In our studies it is clear that cellular vesicular organelles in proximity of the junctions get trapped by the tweezers and they are ultimately the objects pushing the junctions. This causes the trap stiffness to vary between measurements, as it depends on the physical properties of the vesicles. For an absolute measurement of the tensions, we would need to calibrate the trap stiffness for each organelle trapped before each measurement. There are techniques that offer active tweezers calibration in-vivo ^28^, but they rely on forward scattering interferometry, which is challenging, at best, in thick scattering media such as a chick embryo sample. Modification of the system to work in back-scattering mode should be possible for future work ^29 30^. Even without quantitative force measurements we have shown that, despite the variance in stiffness introduced by trapping organelles, we can still measure significant differences in tension of populations of junctions in different areas of the chick embryo. Junctions in the posterior area of the embryo aligned perpendicular to the A-P axis were deformed on average significantly less than junctions aligned in the same direction in the central area of the embryo. Junctions in the posterior mesendoderm aligned perpendicular to the A-P axis, in the direction of tissue contraction, showed significantly higher average tension than junctions of mesoderm cells aligned in perpendicular direction, along the A-P axis. Furthermore, the observation that application of myosin inhibitors resulted in a significant loss of junctional tension is in line with the hypothesis the observed myosin cables are responsible for the differences and alignment of junctional tension in the mesendoderm. These observations also support the hypothesis that myosin generated differential tension drives the observed directional intercalation of mesendoderm cells in the early pre-streak embryos. It remains, however, to be shown directly that local changes in tension can result in local myosin accumulation and induce directed intercalation. This will need perturbation experiments to be performed on much longer time scales and will require suitable in vivo indicators of myosin activity^31^.

In all experiments we observed cases where the junctions did not return to their original rest position during the time frame of the measurements. In line with this we have found that a simple Maxwell viscoelastic model, with only a few parameters fits our experimental data best. This agrees with measurements in the gastrulating Drosophila embryo ^17^. We have used this fitting to obtain the relaxation times with which the junctions relax after release from the trap. We did not observe a significant change in the relaxation time when measuring junctions in different areas of the embryos. There was however an increase in relaxation time after treatment with the myosin I and myosin II inhibitors. This relaxation time is a combination of a time constant deriving from the viscoelastic properties of the junctions themselves, T, and one deriving from the viscosity of the cytosol, 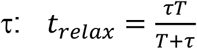. We assume that the properties of the cytosol remain invariant between the different measurements and that the observed differences in relaxation times are due to differences in the junction viscoelastic time constant. The degree of irreversible deformation depends on the dissipation time constant. In Drosophila it has been shown that this dissipation constant is around 60 seconds and that this is highly dependent on the actin cytoskeleton or dynamics ^17^. We deformed the junctions only for a relatively short time, but nevertheless we observed in some cases an irreversible deformation junction. When we looked at the time dependence the effects were more noticeable when myosin activity was inhibited, suggesting that myosin might have role in the setting the dissipation time constant. This will require further characterisation using a greater range of deformation times.

Taken together our findings agree with the hypothesis that in the sickle shaped mesendoderm area the junctions are under higher tension than junctions in other parts of the embryo ^10^. Furthermore they strongly support the notion that this tension difference is due to the accumulation of active myosin II at these junctions. Our experiments further confirm the role of myosin I in the generation of tension in cell junctions in the chick embryo. Future investigations will focus on the determination of the trap stiffness, to obtain absolute value of the junction tensions and on a better characterisation of dissipation characteristics of the junctions in different parts of the embryo and at different time-scales as well as the direct and indirect contribution of various myosin classes on tension generation.

## Methods

### Trapping and Imaging

We performed all experiments reported here with a custom-built inverted fluorescence microscope using an infrared trapping laser ^21^. Fig. S1 shows a diagram of the experimental system. The sample was first imaged with a low-magnification microscope in order to locate the area of interest in the embryo (Fig S1). A custom-made LED ring illuminated the sample with white light at a 45 degrees angle, while images were collected with a 4x microscope objective (Zeiss Achroplan, air immersion, NA=0.1, W.D.=11.1mm) in reflection mode. For fluorescence excitation of the EGFP in the cell membrane, a 100x microscope objective (Nikon CFI Apochromat TIRF, oil immersion, NA=1.49, W.D. = 0.12 mm) working in epifluorescence mode was used in combination with a 488 nm excitation laser (488 Sapphire SP, Coherent). The images were collected onto a scientific CCD camera (Hamamatsu Orca flash 4.0). Excitation and emitted fluorescence light were separated by a dichroic mirror (Chroma ZT488rdc) under the microscope objective and an edge filter (Semrock BLP01-488R-23.3-D) before the camera. The same 100x microscope objective was used to generate the trap by focusing an infrared laser (wavelength 1070nm, ytterbium doped fibre, IPG Photonics). An additional dichroic mirror (Thorlabs DMSP950) and a bandpass filter (Chroma ET750sp-2p8) eliminated the backscattered reflections of the laser onto the camera. The trapping laser was moved using a piezo-actuated mirror (Thorlabs POLARIS-K1S2P) located in a plane conjugate with the back focal plane of the microscope objective. The piezo-actuated mirror and the camera were hardware triggered (National Instruments data acquisition card NI PCI-6251 with connector box SCB-68A) via a Matlab script to ensure that every frame could be assigned a specific trap location. Before every experiment, we removed the bandpass filter and recorded the backscattered reflection of the trap onto the camera to infer the trap position during the experiments. All experiments were performed at 750mW laser power, as measured in the image plane. The trap moved of a total distance of 2.6μm over a time interval of 5 seconds and was kept in the final position, while still on for 2.5 seconds and then turned off for another 2.5 seconds. During the experiment, the embryos were kept at 37 degrees Celsius in a custom-built incubator chamber.

### Embryo sample preparation

A membrane-localized GFP transgenic chicken line was used in all experiments ^10^. Fertilised eggs were obtained from the national Avian Research Facility at the Roslin institute in Edinburgh (http://www.narf.ac.uk/chickens/transgenic.html).

Embryos were incubated in the egg for 5hrs at 37°C by which time in many embryos a well-defined Koller’s Sickle was visible ^32^. The embryos were isolated in EC culture ^33^. For the experiments the embryos were mounted with the epiblast layer facing downwards on a Willco 3cm glass bottom cell culture dish and covered with 2 ml of low viscosity light silicon oil (viscosity 5cSt, Sigma 317667) to prevent drying out of the embryo. The experiments were performed within one hours of mounting the embryos.

Sets of position dependent measurements in the posterior and middle positions were performed on the same embryo. Typically, 10 junctions oriented perpendicular to the AP axis were measured in both positions within an experimental time of 1h.

### Myosin Inhibitors experiments

The myosin inhibitor experiments consisted of sets of measurements before and after addition of the myosin inhibitors. 5h old embryos were mounted in a glass bottom petridish and trapping experiments were performed for 30 min on junctions located in the posterior area of the embryo aligned perpendicularly to the A/P axis. The same embryos were then carefully lifted from the glass bottom and treated with a 10μmolar PBP/PCP solution. After an incubation time of 20min, the trapping experiments were repeated in the posterior area of the sample.

### Data processing and data analysis

A custom MATLAB GUI was implemented to control the components of the set-up and to trigger the camera. The images collected through MATLAB were saved as archival format video (mj2), while a.mat file stored the instrument settings of the measurements. For the analysis, we applied a Gaussian smoothing filter and stretched the contrast for each frame. Through a custom MATLAB function, we produced kymographs for every location along the junction, generating a stack of kymographs. For each kymograph, we identified the location of the junction: we observed that performing a Gaussian fit of the fluorescence intensity along the kymograph lines to achieve subpixel resolution was not effective for our data and failed to identify the junction position because of the low signal to noise level and the presence of scattered light from the organelles in proximity of the junction. Therefore, we adopted a modified version of a seam carving algorithm. Seam carving algorithms are designed for content-aware resizing ^34^ by removing the paths in an image which have minimal variation. In a greyscale image, these paths correspond to the shortest paths between the first row and last row of the image weighting pixel values on the grayscale. Our modified seam carving algorithm finds the shortest path for each kymograph in the stack weighting both on the grayscale value of each pixel in the current kymograph and on the neighbour pixel of the previous and subsequent kymograph in the stack. This approach enhances the precision in determining the junction location at each frame for our datasets. Fig. S3 shows a comparison between using Gaussian fittings and adopting the seam carving algorithm to determine the junction locations. Finally, we identified the kymograph associated with the highest deformation of the junction, we converted the results in *μm* and *s* and we extracted the value of the maximum deformation. By plotting the deformation of the junction against time, we concluded our analysis by fitting the data with the *fit* function in Matlab (with the *Robust mode* on) using as “*fittype*” the equation derived from the Maxwell viscoelastic model. The data were aggregated and compared. Significance value were obtained by performing significance tests on the data sets to reject the null hypothesis (with Matlab *ranksum* function).

### Viscoelastic Model

To describe the dynamics of cell-cell junctions, we used a Maxwell viscoelastic model. This model describes the junction as if it were a purely elastic element of elastic constant *E* and a purely damping element of damping coefficient *η* (Fig. S7).

When the junction is deformed by a strain *x*, it is subjected to a tension force F_j_ opposing the deformation:

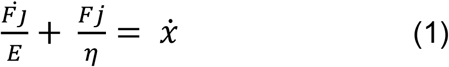

During the optical manipulation, the balance of forces reads as:

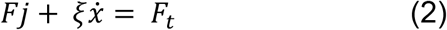

Where *ξẋ* is the drag in the cytosol for an object (the manipulated junction) moving with speed *ẋ, F*_*t*_ is the optical force.

Since we could not characterise the stiffness of the optical tweezers, we used the model to fit the relaxation time after switching off the trapping laser. When the tweezers are off, the tension is balanced only by the drag in the cytosol opposing the return of the junction to the rest position:

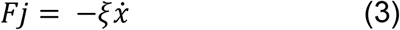

Using this definition of the force with Eq.1, we can derive the differential equation:

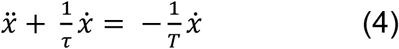

Where *τ* = ^*η*^/_*E*_ represents the relaxation time of the junction and *T* = ^*ξ*^/_*E*_ is the relaxation time of the cytosol.

The solution that describes how the junction moves back to its rest position is a single exponential curve:

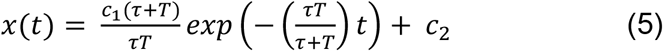

Where c_1,2_ are obtained by applying the initial conditions, i.e. the position x_m_ of the junction at t=0 and its velocity vm at t=0+:

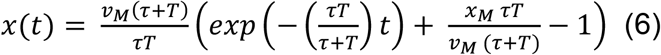

We used the *fit* function in Matlab with *Robust mode on*. We fitted the data with a solid linear solid model and with a Kelvin-Voigt model in addition to the described Maxwell model, but we found that the Maxwell model outperformed the other fits for accuracy (Fig. S7).

## Acknowledgments

This project has received funding from the European Union’s Seventh Framework Programme for research, technological development and demonstration under grant agreement no 608133 and grant BB/N009789/1 from the BBSRC. Funding was also received from the Scottish Universities Physics Alliance (SUPA).

---

**Figure S1.**
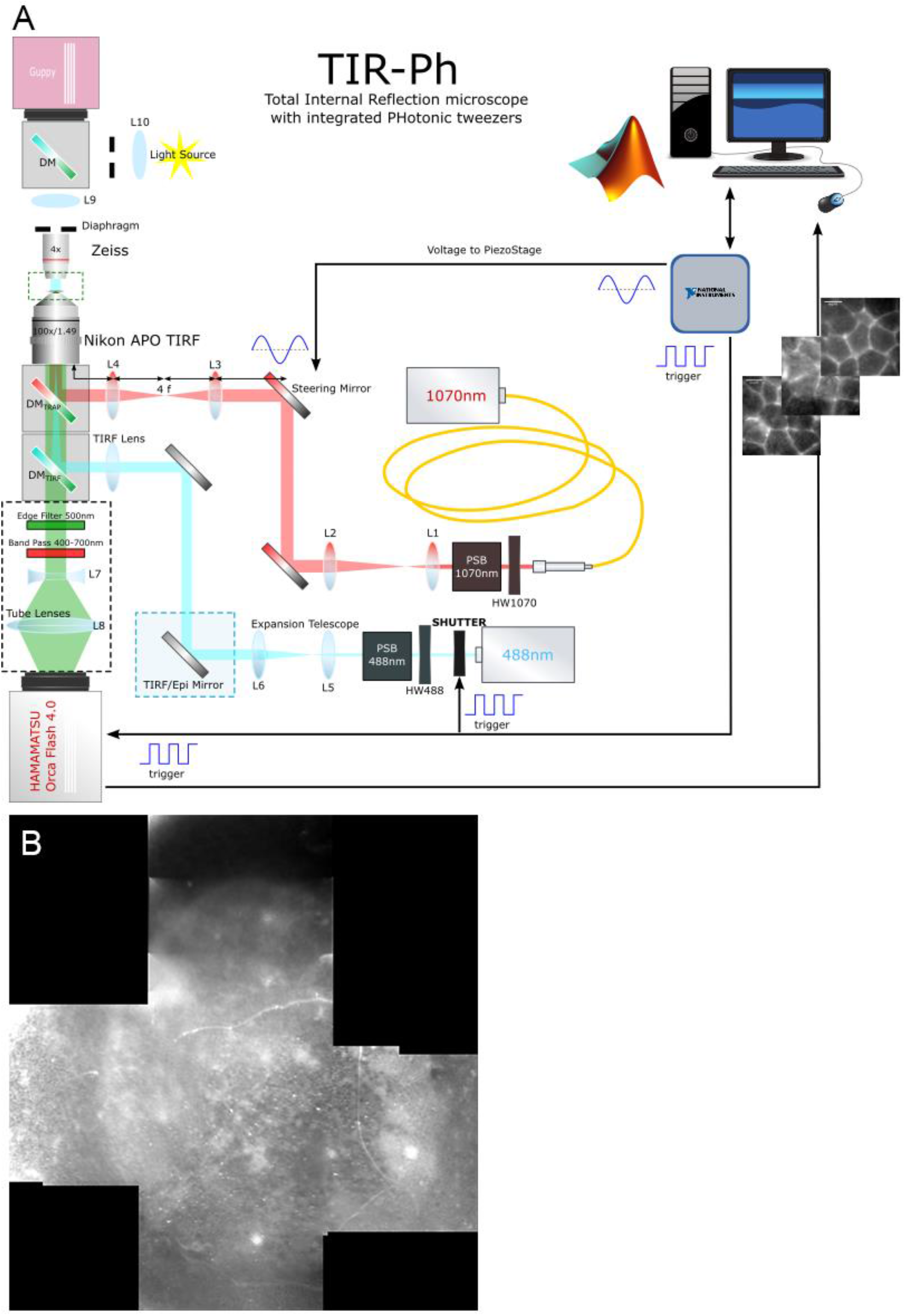
(A) Optical setup combining epifluorescence microscopy and optical tweezers. For details of setup see materials and methods (B) Example of low magnification image of the chick embryo taken with the Guppy camera. Low magnification images were used to orientate in the embryo.

**Figure S2.**
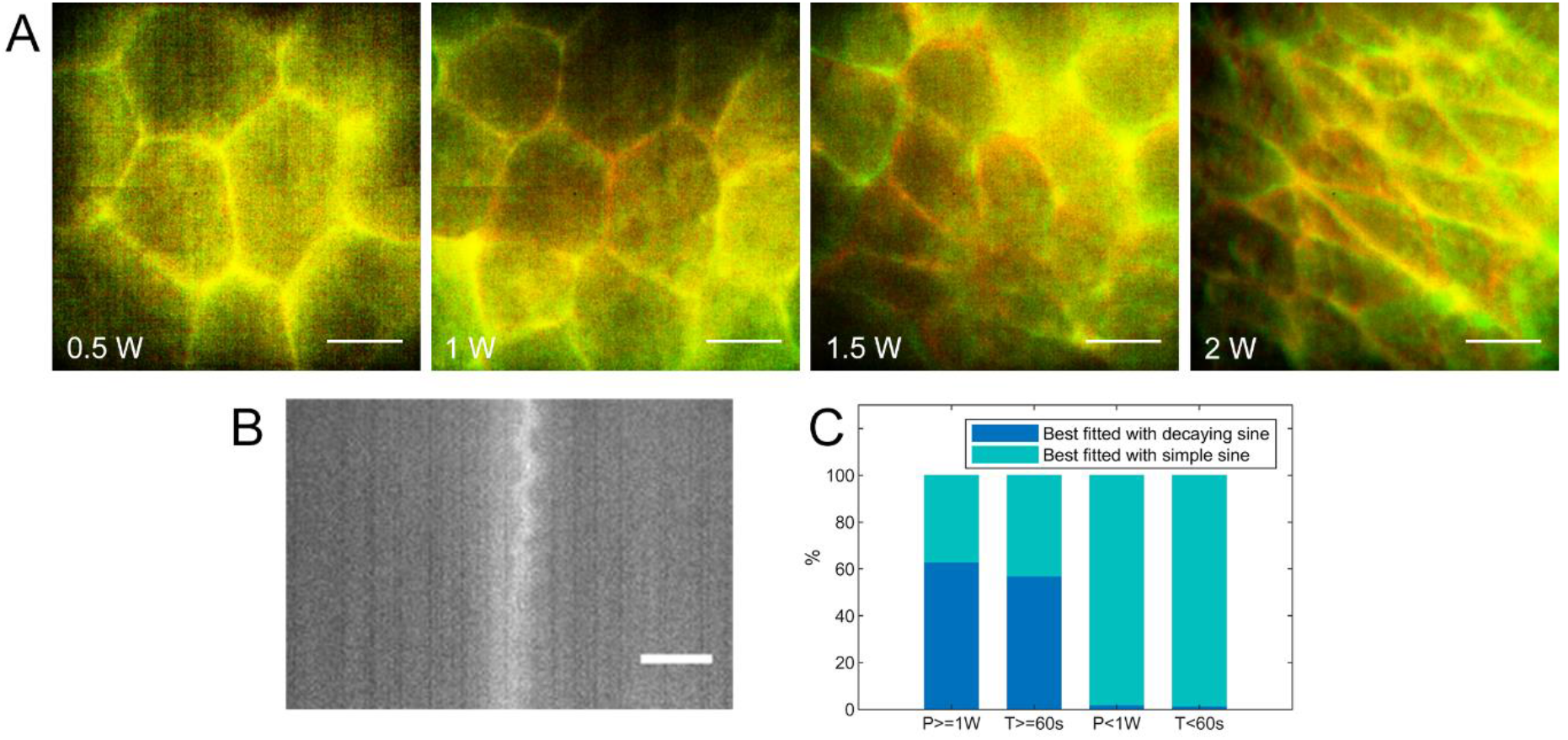
Power damage on chick embryos junctions. (A) False colour images of a chick embryo subjected to different laser powers: in red the embryo at time t=0, before the laser is turned on; in green the same embryo observed after the laser was left on for 30 seconds. For laser powers <1 W, there is no visible damage: the first frame and the last frame overlap. For power = 1 W, there is minimal damage. At powers >1.5W, major damage is observed: the junctions in the embryo relax and a long range deformation is observed. (B) shows the kymograph of a junction being subjected to sinusoidal motion of the trapping laser (see Fig. S4). The laser power was 1W and the duration of the measurement was 60 seconds. The kymograph shows that the junction follows the trap movement initially, but the amplitude of deformation reduces over time. This behaviour can be fitted by using an exponentially decaying sine function. We compared junctions subjected to sinusoidal motion with different parameters (power <1 W, power >= 1W, duration <60s, duration >=60s) and verified whether they would be best-fitted by a decaying sine wave (C). We observed that the decay behaviour was predominantly observed after longer exposure to the laser and at higher powers. This suggests that the junctions change during long exposure and or higher power. Based on these findings we chose a constant power of 750mW. At this power the embryos showed no visible damage, while we were able to move a higher percentage of junctions than by using lower power (i.e. 500mW).

**Figure S3:**
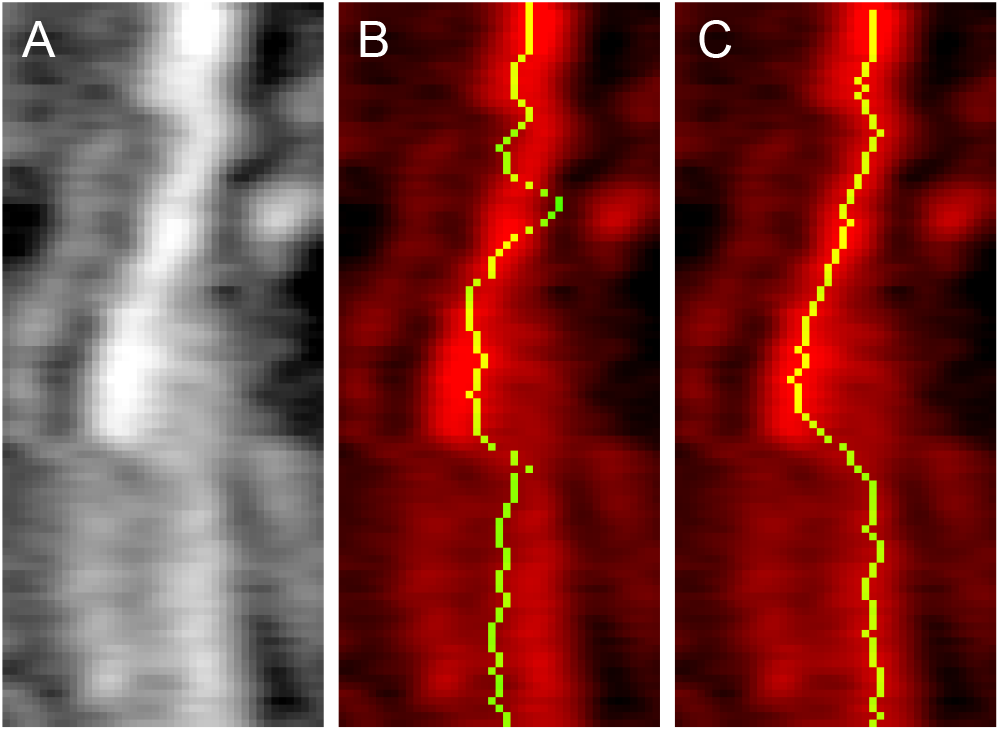
Gaussian fit vs seam carving algorithm comparison. (A) shows a junction kymograph. (B-C) False colour images with the derived junction positions overlaid in yellow over original kymograph: in (B) the junction positions were extracted by fitting each line profile of the kymograph with a Gaussian curve; in (C) the junction positions were extracted by using an implementation of the seam carving algorithm. The sequence (A-C) shows that the seam carving algorithm better reproduces the position of the junction that that obtained by using a Gaussian fit.

**Figure S4.**
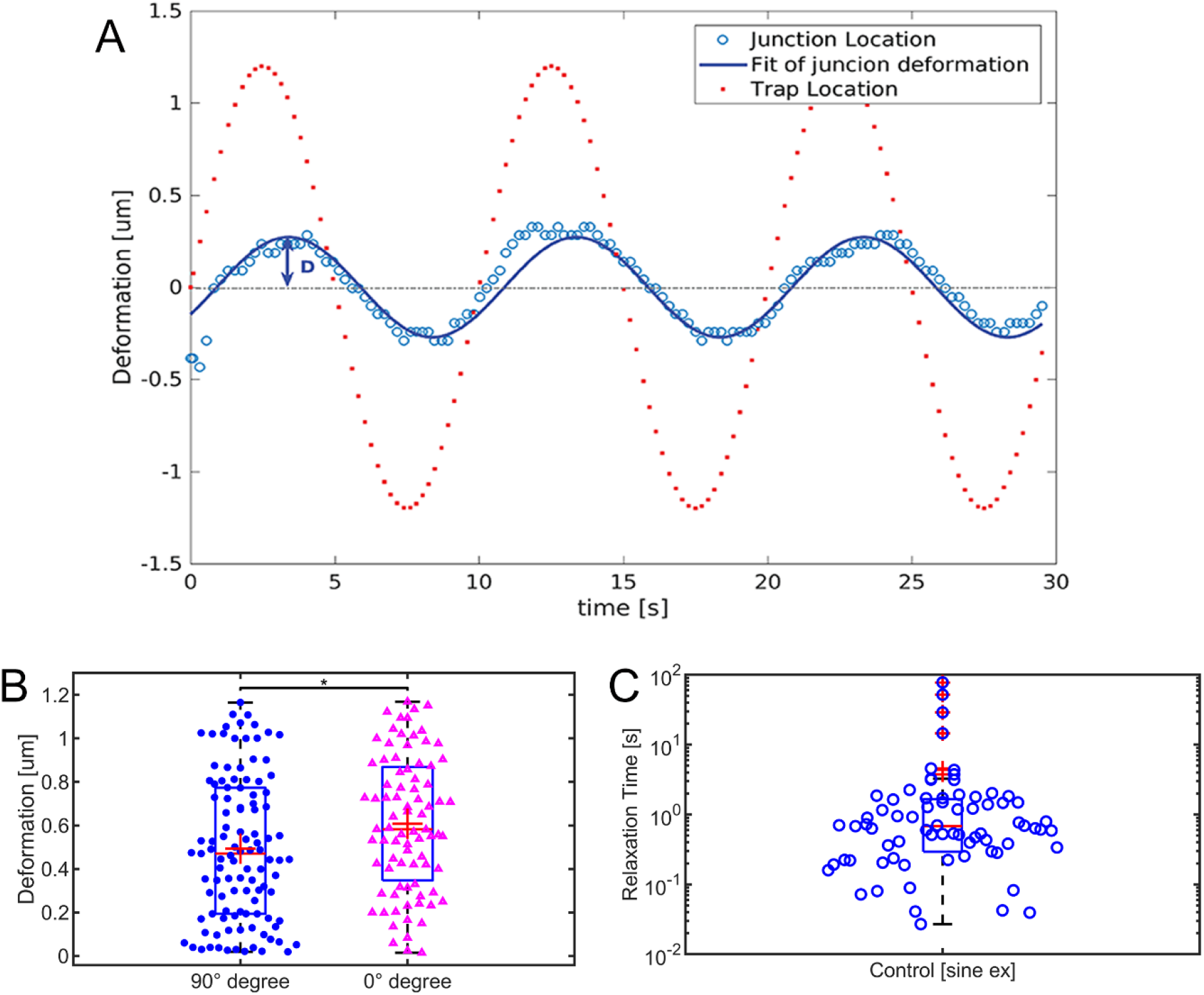
Deformation of junctions perpendicular and parallel to the A/P axis. (A) Example of position of the junction over time (blue empty circles) with its fit (blue line) compared with the position of the trapping laser (red dotted line) for sine wave pulling experiments. In these experiment the trapping laser moved back and forth with a sinusoidal motion rather than the pull and release approach used in the majority of the experiments described in the main text. The trap oscillated with a sinusoidal motion amplitude of 2.6 μm and a frequency of 0.1 Hz for a duration of 30s. We extracted the movement of the junction over time by applying the seam carving algorithm to the kymograph of the junctions. The position of the junction over time was fitted with sine function and we extracted amplitude, phase and period. (B) Boxplot and distribution of the maximum deformation of junctions as fitted from the sine wave experiments in cell junctions with different alignment respect to the A-P axis: junction perpendicular to the A/P axis, and therefore aligned to the super-cellular Myosin cables (blue filled circle, n=109 collected over 4 embryos) had a median deformation of 0.45 μm, while junctions parallel to the A/P axis (pink empty triangles, n=84 collected over 4 embryos) had a median deformation of 0.6μm. (C) From the phase shift between the trap and the junction movement, it is possible to estimate the relaxation time of the system. According to viscoelastic models, t_relax_ = tan(φ), where φ is the phase shift expressed in degrees. We measured a relaxation time of 0.68s for junctions perpendicular to the A/P axis, which is in agreement with the values measured by using the viscoelastic fitting in the pull & release experiments.

**Figure S5:**
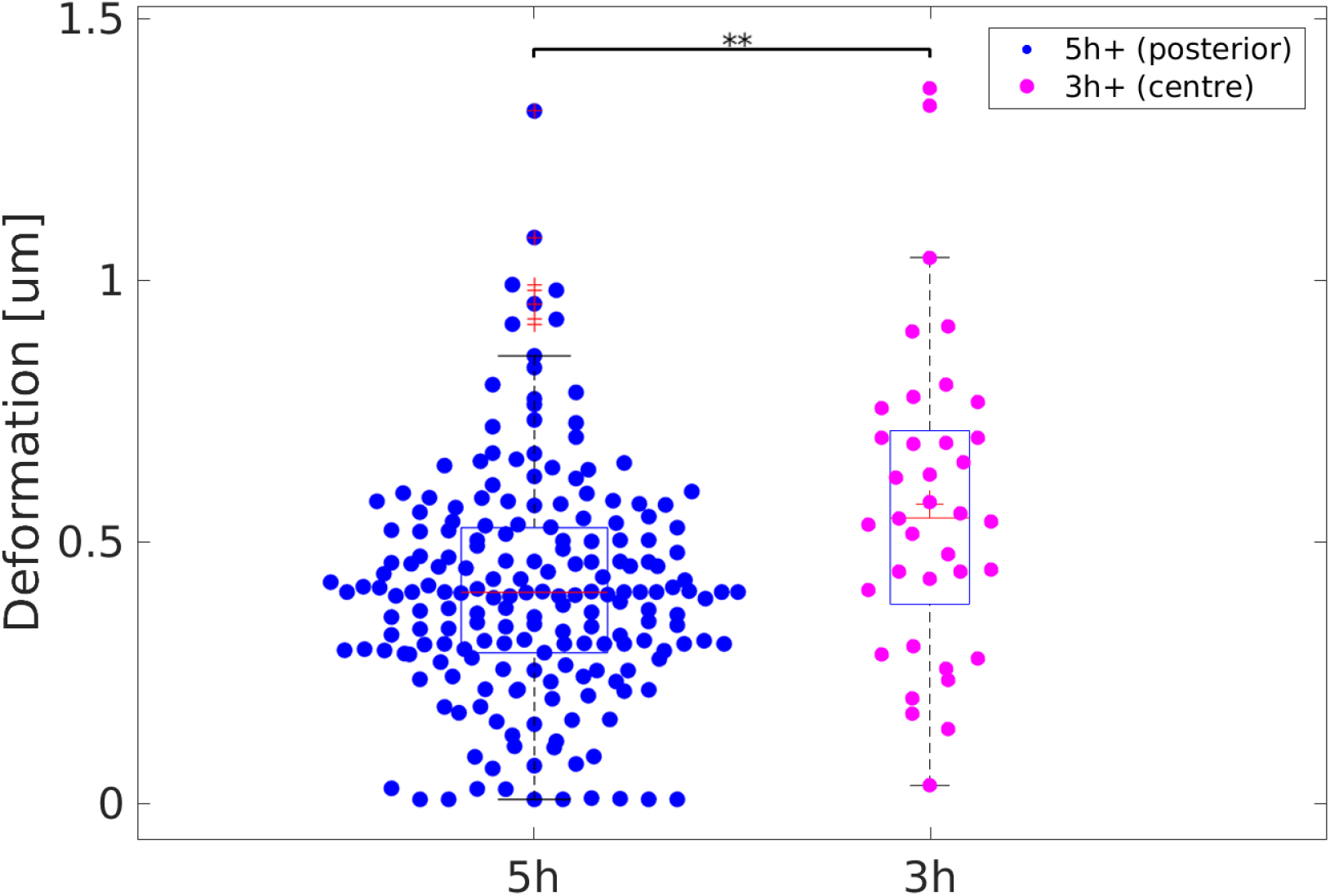
Deformation of junctions in chick embryos of different age. Boxplot and distribution of the maximum deformation of junctions measured in the posterior of 5h old embryos (blue filled circle, n=203) and in the centre of 3h old embryos (pink circles, n=37). Median values for the deformation are 0.39μm 0.59μm for the 5h old and the 3h old embryos respectively. The data for the 5h old embryos were aggregated from ten different embryos (Control embryos in the main text), while the data for the 3h old embryos were aggregated from two different embryos each. ** indicates p-values <0.01. The significant difference between the two dataset reinforces the hypothesis that junctional tension increases with the presence of super-cellular Myosin II chains.

**S6.**
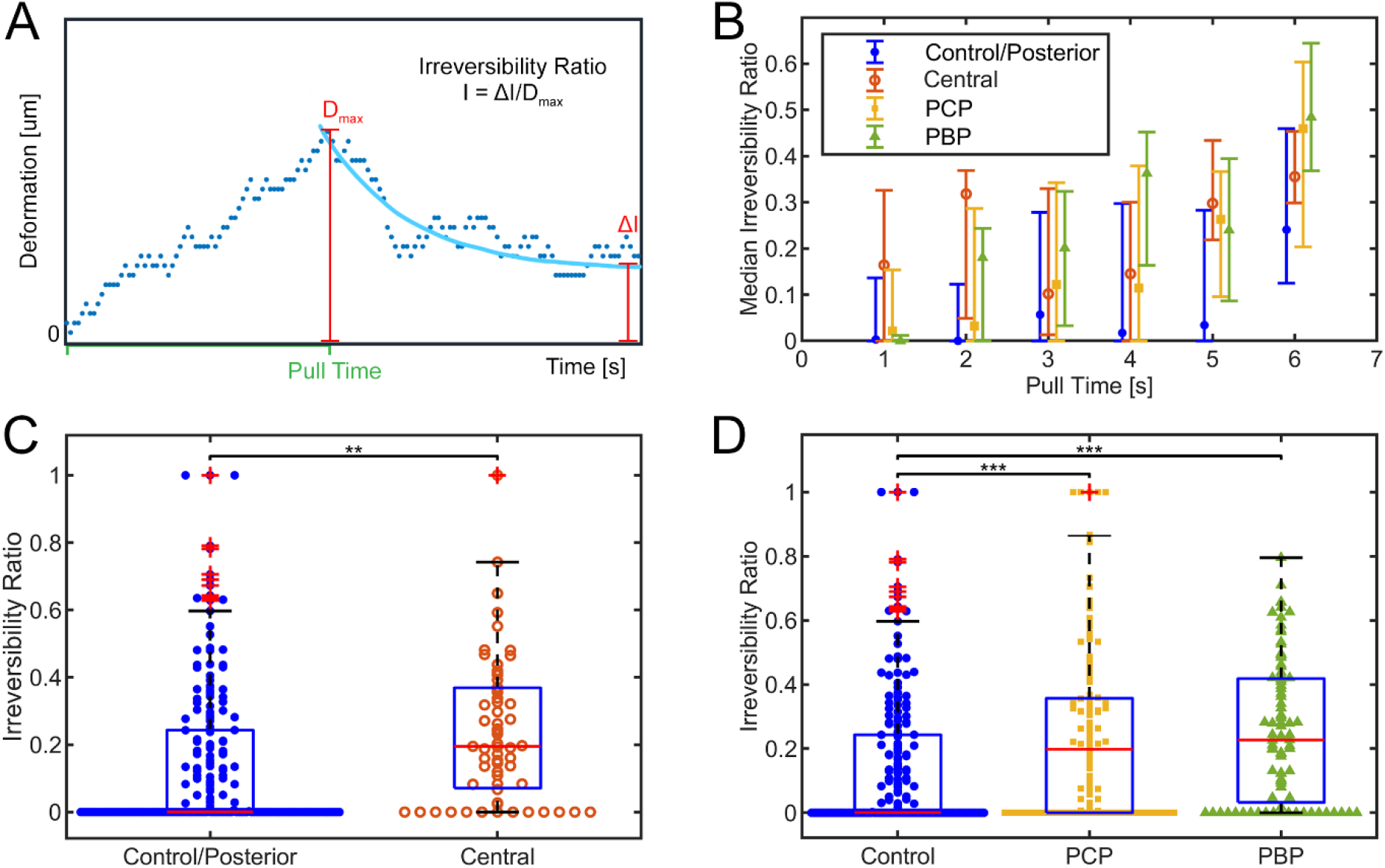
Irreversibility Ratio. (A) Cartoon of the definition of irreversibility ratio: the ratio between the position of the junction after being pulled at infinite time ΔI (as measured through the Maxwell model) and the maximum deformation of the junction Dmax (red lines), (B) Median Irreversibility ratio as function of pulling time. The data are aggregated in intervals of pulling time of 1 second. Pull times are defined as the time which the junction is released from the tweezers (Green line in A). Error bars represent the 25 and 75 percentiles. (C-D) Boxplot and distribution of the irreversibility ratio of controls (blue filled circles, n=203) compared with that of junctions measured in the central area of the embryo (C, empty circles, n=57) and compared with that measured in embryos treated with Myosin inhibitors PCP (D, yellow squares, n=132) or PBP (green triangles, n=88. ** indicates p-values <0.01; *** indicates p-values < 0.001.

**S7.**
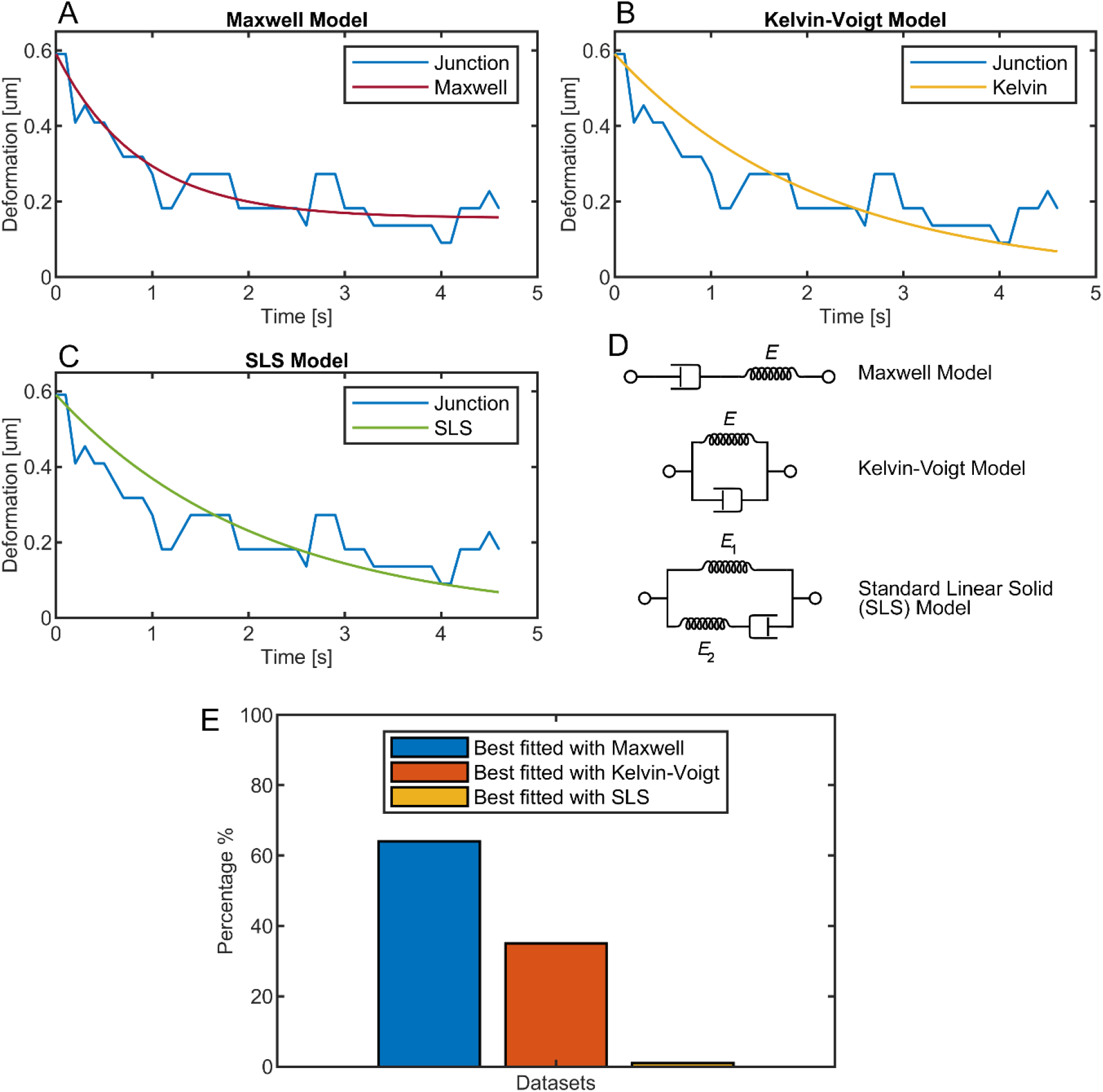
Comparison of goodness of fit of three visco-elastic models. (A-B) Example of a junctional relaxation kinetics (blue line) fitted with Maxwell model (A), with Kelvin-Voigt model (B) and with SLS model (C). To decide the best fit, we compared two parameters, the adjusted R-squared and the absolute error on the measured variables. For the junction in the example, the Maxwell fitting presented an Adjusted R-squared of 0.84 and absolute error on fitting variable of 0.07; the Kelvin-Voigt fitting presented an R-squared of 0.38 and absolute error on fitting variable of 0.57; the SLS fitting presented an R-squared of 0.35 and absolute error on fitting variable of 1E16. We concluded that the best fit was obtained by using the Maxwell model fitting. (D) Schematic representation of the three models compared. (E) By fitting our datasets for the push & release experiments with all three models, we observed that the Maxwell model best fits most of our data. The major difference between the Maxwell and the Kelvin-Voigt model is that the first accommodates the irreversibility of the junction deformation, while the second provides a fitting that will always return to the rest position at 0. The data are the based on analysis of 203 junctions in control experiments of various embryos.

## Movie legends

**Movie 1:** Example of Pull & Release experiment showing the deformation of a junction. The video was acquired at a frame rate of 16.4 frames / second, and it is reproduced at 16 frames / second. Scale bar is 5 um.

**Movie 2:** Example of Pull & Release experiment showing the deformation of a junction when pushed by an organelle. The video was acquired at a frame rate of 10 frames / second. The video rate is 14 frames / second. Scale bar is 3 um.

